# Structural preferences shape the entropic force of disordered protein ensembles

**DOI:** 10.1101/2023.01.20.524980

**Authors:** Feng Yu, Shahar Sukenik

## Abstract

Intrinsically disordered protein regions (IDRs) make up over 30% of the human proteome and instead of a native, well-folded structure exist in a dynamic conformational ensemble. Tethering IDRs to a surface (for example, the surface of a well-folded region of the same protein) can reduce the number of accessible conformations in IDR ensembles. This reduces the ensemble’s conformational entropy, generating an effective entropic force that pulls away from the point of tethering. Recent experimental work has shown that this entropic force causes measurable, physiologically relevant changes to protein function, but how the magnitude of this force depends on the IDR sequence remains unexplored. Here we use all-atom simulations to analyze how structural preferences encoded in dozens of IDR ensembles contribute to the entropic force they exert upon tethering. We show that sequence-encoded structural preferences play an important role in determining the magnitude of this force and that compact, spherical ensembles generate an entropic force that can be several times higher than more extended ensembles. We further show that changes in the surrounding solution’s chemistry can modulate IDR entropic force strength. We propose that the entropic force is a sequence-dependent, environmentally tunable property of terminal IDR sequences.

## Introduction

Intrinsically disordered proteins and protein regions (IDRs) do not have a native, stable structure. Instead, IDRs exist in a constantly interchanging conformational ensemble that contains transient and relatively weak intramolecular interactions. These interactions define the structural preferences and the resulting average shape of the ensemble. Decades of work have linked the structural preferences of IDRs to their biological functions^1–4^.

Like other polymers, IDR ensembles have a high conformational entropy. This conformational entropy can be reduced by covalently linking, or tethering, the IDR through one of its termini to a surface (**Fig. 1A**). In this case, entropy is reduced due to the constraint placed upon the ensemble by the surface it is tethered to. As a result, upon tethering a polymer chain will try to maximize its conformational entropy by producing an effective force that pulls up and away from the point of tethering, gaining entropy by increasing its number of accessible conformations (**Fig. 1B**). For homopolymers, the strength of this entropic force is determined by the polymer’s length and the geometry of the constraining surface^5^ as shown both theoretically and experimentally^6,7^.

**Figure 1.**
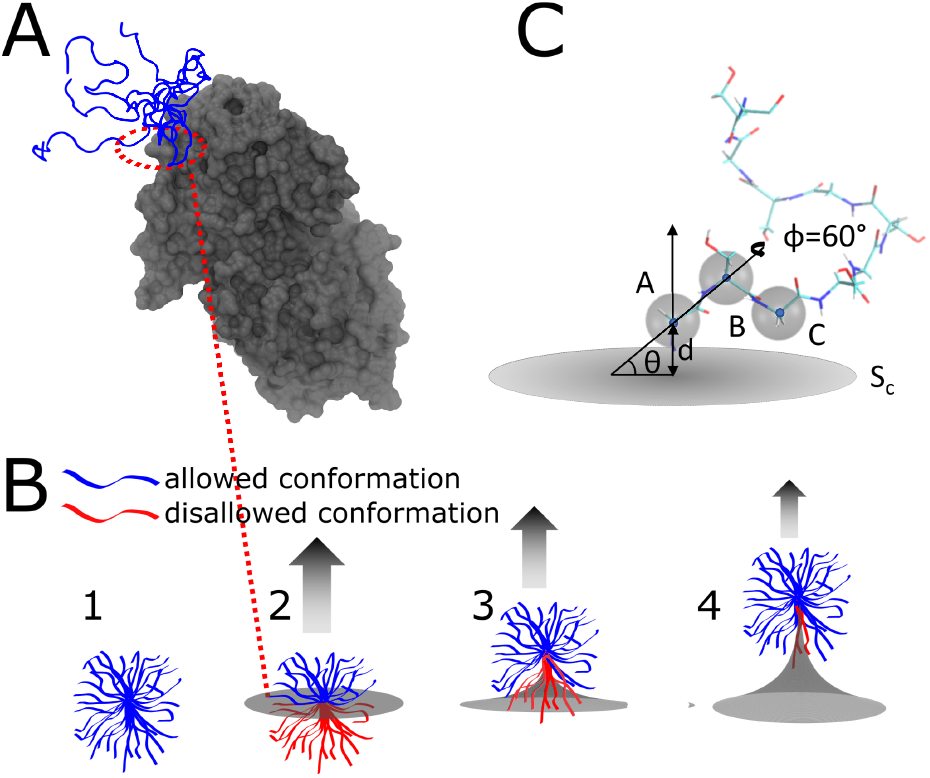
Dynamic IDR conformational ensemble generates an entropic force. **(A)** IDR tethered to a well-folded domain. Here, the C-terminal IDR tail of the UDP-glucose 6-dehydrogenase (UGDH) protein is shown in blue (with 5 overlapping conformations to illustrate the variability in the ensemble) tethered to the main folded domain of the enzyme (in grey)^14^. (**B**) Schematic showing how a constraining surface alters the conformational entropy of an IDR ensemble. (1) A few representative conformations from an IDR ensemble (blue) occupy an extended volume. (2) as the ensemble is tethered at the terminal to a surface (grey), some conformations clash with the surface (colored in red), causing them to be disallowed and lowering the conformational entropy. (3 and 4) The number of accessible tethered states (Ω_*T*_) can be regained by “pulling up” against and pinching the surface (arrow). The ratio between the allowed and total number of conformations for a given ensemble is proportional to the entropic force strength (see **Eq. 3**). (**C**) Enhanced conformational sampling. All conformations of an IDR are aligned along the vector AB connecting the first two *C*_α_ atoms. The distance *d* between the constraining surface S_c_ and point A is varied to represent tether flexibility. The angle between vector AB and the constraint surface, θ, is varied to represent one degree of rotation for the ensemble, and a second angle, ϕ, represents the rotation angle along the AB vector.

This tethering scenario may seem rare when considering naturally occurring proteins, but it is actually rather common: in eukaryotes, IDRs are often tethered to a more rigid surface that constrains the chain’s conformational entropy, and this tethering results in measurable effects. For example, IDRs tethered to a cell membrane can sense the curvature of the membrane and help to facilitate the endocytosis process through entropic force^8–11^. The same entropic force can also help translocate IDRs through the bacterial cell wall to the extracellular environment, an essential process for bacterial infection^12,13^. An even more prevalent scenario occurs when disordered N-or C-terminal IDRs are attached to a well-folded protein region (**Fig. 1A**). The entropic force exerted by such disordered terminal regions can influence protein function, including ligand binding affinity and thermodynamic stability^14,15^. These examples suggest that entropic force may be an important and prevalent mechanism unique to IDRs that mediates biological function.

The entropic force literature primarily focuses on the role of an IDR sequence’s length. IDR length is indeed a critical factor in determining entropic force magnitude^9,14^, since the longer the chain, the higher the number of conformations available. But is chain length always the most dominant factor affecting entropic force magnitude? Previous research has shown that, unlike homopolymers, IDR ensembles have distinct sequence-encoded structural preferences^16–24^. These structural preferences affect the average shape occupied by IDR ensembles^25,26^. We hypothesize that besides the length, the sequence-encoded shape of the ensemble will also determine the entropic force exerted by tethered IDRs.

To test this hypothesis, we use all-atom Monte Carlo simulations to sample the conformational ensembles of over 90 experimentally validated IDR sequences. To gauge the magnitude of the entropic force sequences can exert, we measure the reduction in the number of allowed conformations upon tethering their ensembles to a flat surface. Our simulations show that the entropic force depends not only on the length of the IDR but also on its sequence-encoded ensemble shape, with more compact ensembles exerting a stronger entropic force. To further test this finding, we alter the dimensions of each ensemble by changing their interaction with the surrounding solution (while keeping the sequence intact). We show that solution-induced compaction also increases the entropic force, but only for a subset of the sequences. Our findings reveal how sequence-encoded intramolecular and protein:solution interactions combine to modulate the magnitude of the entropic force exerted by tethered IDR. They also suggest that the entropic force can be tuned by evolution to exert an optimized effect on full-length proteins.

## Materials and Methods

### Intrinsically disordered protein prediction with AlphaFold database

Systematic evaluations of AlphaFold2 (AF2) previously showed that it is a good predictor of intrinsically disordered regions^27–29^. We downloaded the predicted structures of three different proteomes (Saccharomyces cerevisiae: UP000002311, Arabidopsis thaliana: UP000006548, Homo sapiens: UP000005640) from the AF2 database version 3.^30^ The disorder predictions are obtained from AF2’s pLDDT score. Based on a previous report^27^, we used 30 consecutive residues with pLDDT < 50% as an indicator for IDRs. Detected IDRs are labeled as terminal if they start at the N-terminal or end at the C-terminal of the protein in the AF2 database.

### All-atom Monte-Carlo simulation

All IDRs were simulated with the ABSINTH implicit solvent force field using the CAMPARI simulation suite v2_09052017.^31^ The example parameter file and simulation settings are provided in the GitHub repository. Simulations were conducted at 310 K with 10^7^ steps of equilibration. After calibration, production conformations were written every 12,500 steps. For each IDR, we performed five independent simulations with ∼5,600 individual conformations in each repeat. This leads to a total of ∼28,000 conformations for each IDR (details in **Table S1**). For PUMA scrambles, we performed three individual repeats, leading to ∼16,800 conformations.

### Calculation of ensemble properties

#### Normalized end-to-end distance

The end-to-end distance *R*_*ee*_ of polymers can be calculated based on the number of residues in the chain (N) using 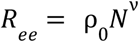. The scaling law ν can have a range of fractional values. Specifically, ν = 0. 59 for expanded chains, ν = 0. 5 for ideal (or θ-state) polymers, and ν = 0. 33 for collapsed/compacted polymers. For homopolymers, the prefactor ρ_0_ is constant and depends on the segment length of the monomer^23,32^. Seven glycine-serine dipeptide repeat (GS-repeats) sequences with 8, 16, 24, 32, 40, 48, and 64 GS segments were simulated and analyzed as described above with five individual repeats. The GS-repeat *R*_*ee*_ data were fitted to a power-law function based on the equation above (**Fig. S1**). The fitted exponent for GS-repeats is ν = 0. 48 ± 0. 03 demonstrating ideal polymer behavior, in agreement with previous experimental data^33,34^. We use this fitted curve to normalize IDR *R*_*ee*_ for comparison across IDRs of different lengths. We interpolate and extrapolate corresponding GS-repeats *R*_*ee*_ based on the length of the IDR of interest. We calculate the normalized *R*_*ee*_ using the following equation.

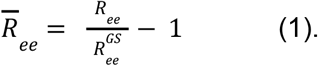

Here, 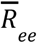 is the normalized end-to-end distance, *R*_*ee*_ is the end-to-end distance of the target IDR, and 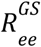 is the calculated end-to-end distance of a GS-repeat sequence of the same length as the target IDR (obtained from the fit shown in **Fig. S1**).

#### Asphericity

IDR ensemble properties were analyzed using the MDtraj python library^35^. *R*_*ee*_ was calculated between the *C*_α_ of the first and last residue of the IDR. Helicity was calculated using the DSSP algorithm integrated into MDtraj^36^. Asphericity was calculated using the gyration tensor of the simulated IDR ensemble as described previously^37–39^.

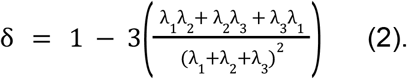

Here, δ is the asphericity, and λ_1, 2, 3_ are the three principal moments of the gyration tensor.

The standard deviation of all these properties was calculated based on the averages from the five independent repeats. Analysis scripts are available at the accompanying GitHub repository at https://github.com/sukeniklab/Entropic_Force.

#### Entropy analysis

To calculate the effect of tethering on IDR conformational entropy we count the number of allowed conformations in the ensemble upon tethering (**Fig. 1B**). To do this, we first tether each conformation of each simulated IDR ensemble to a single point on a flat surface and then calculate the number of allowed conformations Ω_*T*_ from the total number of conformations in the simulated ensemble Ω_*U*_. Tethering is done relative to the first, second, and third *C*_α_ coordinates of each conformation, labeled here as A, B, C (**Fig. 1C**). For each conformation, we move A to the origin of the coordinate system. We plot the constraint surface, *S*_*c*_, perpendicular to the surface containing atoms A, B, and C.

#### Enhanced sampling

In order to better understand the spatial relationship between the ensemble and the constraining surface, we perform several geometric transformations on each sampled conformation for calculating Ω_*T*_ : (1) To account for the possibility of stretching at the point of tethering, we vary the distance *d* between point A and *S*_*c*_. (2) To account for the possibility of rotation around the point of tethering, we vary the half-angle θ formed between the norm vector to *S*_*c*_ with vector AB. (3) We rotate the vector AB with an angle ϕ.

All the coordinates specified here are illustrated in **Fig. 1C**. In total, we make 36 transformations (3 values for *d*, 3 values for θ, and 6 values for ϕ) for each conformation of each simulated ensemble.

#### Entropy calculation

We consider the interaction between IDR and the constraint surface as a hard sphere interaction. Accessible conformations are defined as those with no *C*_α_ that is positioned below the constraining surface. We use the dot product between the norm vector of *S*_*c*_ and the coordinate of *C*_α_ to calculate and determine the relative position of the *C*_α_ to the surface *S*_*c*_. We then count the number of all accessible conformations in the tethered, original ensemble and all *d*, θ, and ϕ permutations. Finally, we sum the number of accessible states from these perturbations and calculate the entropic force strength. The entropic force is then given by

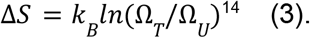

Here *k*_*B*_ is the Boltzmann constant, Ω_*T*_ is the total number of possible IDR conformations when the ensemble is tethered to a surface and Ω_*U*_ is the total number of conformations sampled for the same IDR ensemble when untethered. The entropic force strength is proportional to the Δ*S*. The transformation and analysis scripts are provided as Jupyter notebooks at https://github.com/sukeniklab/Entropic_Force.

#### Ensemble XZ-projections

For each IDR conformation, we move A to the origin of the coordinate system and rotate the conformation to make AB fall on the Z-axis (Z>0).

XZ-coordinate of each *C*_α_ will provide an ensemble projection of IDR ensemble on the XZ plane. The *C*_α_ density was normalized by the number of amino acids in the sequence, the frame number of trajectories, and the bin size.

#### Solution Space Scanning simulations

Solution space scanning simulations are conducted as described previously^16,31,40^. Briefly, we modify the effective Hamiltonian of the ABSINTH force field to alter protein backbone:solvent interactions. The ABSINTH hamiltonian is a sum of four energy terms:

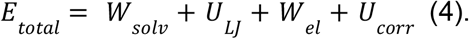

*U*_*LJ*_, *W*_*el*_, and *U*_*corr*_ represent Lennard-Jones (LJ) potential, electrostatic interaction, and torsional correction terms for dihedral angles. *W*_*solv*_ is the solvation free-energy and equal to the transfer free energy between vacuum and diluted aqueous solution. Changing the free energy term *W*_*solv*_ such that results in a change in the protein:solvent relative interaction strength, defined by

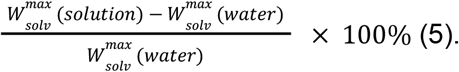

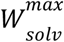 is the solvation free-energy calculated based on fully extended protein conformation in different solution conditions. Negative values of protein:solvent interaction represent solutions that are attractive to the protein backbone, such as urea solutions, while positive values represent solutions that are repulsive to the protein backbone, such as those containing protective osmolytes. A value of 0 represents a buffered, aqueous solution with no cosolutes. We simulated seven different solution conditions for each IDR with a protein:solvent relative interaction strength ranging from +3% (equivalent roughly to 1 M TMAO) to -3% (equivalent roughly to 1.5 M Urea)^40^. It is important to note, however, that even the most attractive solutions used here are not sufficient to unfold well-folded protein domains. We use the same temperature and sampling method for each solution condition as we do for aqueous solutions. The simulation averages of ensemble properties and entropic force in all solution conditions, as well as sequence details, are reported for all IDRs in **Table S1**.

#### Limitations and drawbacks of entropic force calculations

In our calculations, we completely neglect any interactions between the IDR and the surface other than steric, hard-core repulsions. We also assume that the constraining surface is completely flat. In the context of an actual, full-length protein, constraining surfaces will have distinct chemical moieties, including hydrophobic, polar, and charged residues. Specific surface chemistries will introduce an enthalpic component to the free energy change upon IDR tethering which can alter, and sometimes completely reverse, the force induced by tethering. These effects are very important as shown in several cases, especially when charges are introduced^9,41,42^.

Another limitation is that the constraint surface we use is fixed, flat, and does not change over time. The surface of folded domains displays irregular shapes and fluctuations and motions that may change the number of allowed conformations or change the overall entropy of the entire system which we didn’t consider here. Indeed, some of the solution chemistry changes we use in this work may also act to alter these fluctuations.

To mitigate these limitations, we stress that the entire dataset was obtained using the same methods and analysis, and compared against the same GS-repeat benchmarks. This self-consistency is what allows us to probe the role of the ensemble itself on the entropic force, all other factors being held constant.

## Results and Discussion

### The human proteome is rich in disordered terminal sequences

We define terminal IDRs as those that exist at the N or C termini of proteins, and reason that with one free end, such IDRs can exert an entropic force against the more rigid, folded protein domain to which they are connected (**Fig. 1A**). To see if terminal IDRs are common in proteomes, we tested their prevalence in the yeast, arabidopsis, and human proteomes. using the AlphaFold Protein Structure Database v3^43^ (**Fig. 2A**). The confidence score of AlphaFold2 (pLDDT) has been previously shown to be a good indicator of potential disordered regions^27^ and so was used to identify disordered regions in the three proteomes. A protein segment was marked as disordered when it had more than 30 consecutive residues with a ‘very low’ pLDDT score (< 50%). For the proteomes we tested, over 40% of proteins have at least one disordered segment, in line with previous studies^44^ (**Fig. 2A**, left). In the human proteome specifically, over half of the proteins that contain IDRs have at least one at either the N-or C terminal (**Fig. 2A**, right). This result indicates that terminal-tethered IDRs exist widely in eukaryotes and that the entropic force scenario described above can occur in many proteins.

**Figure 2.**
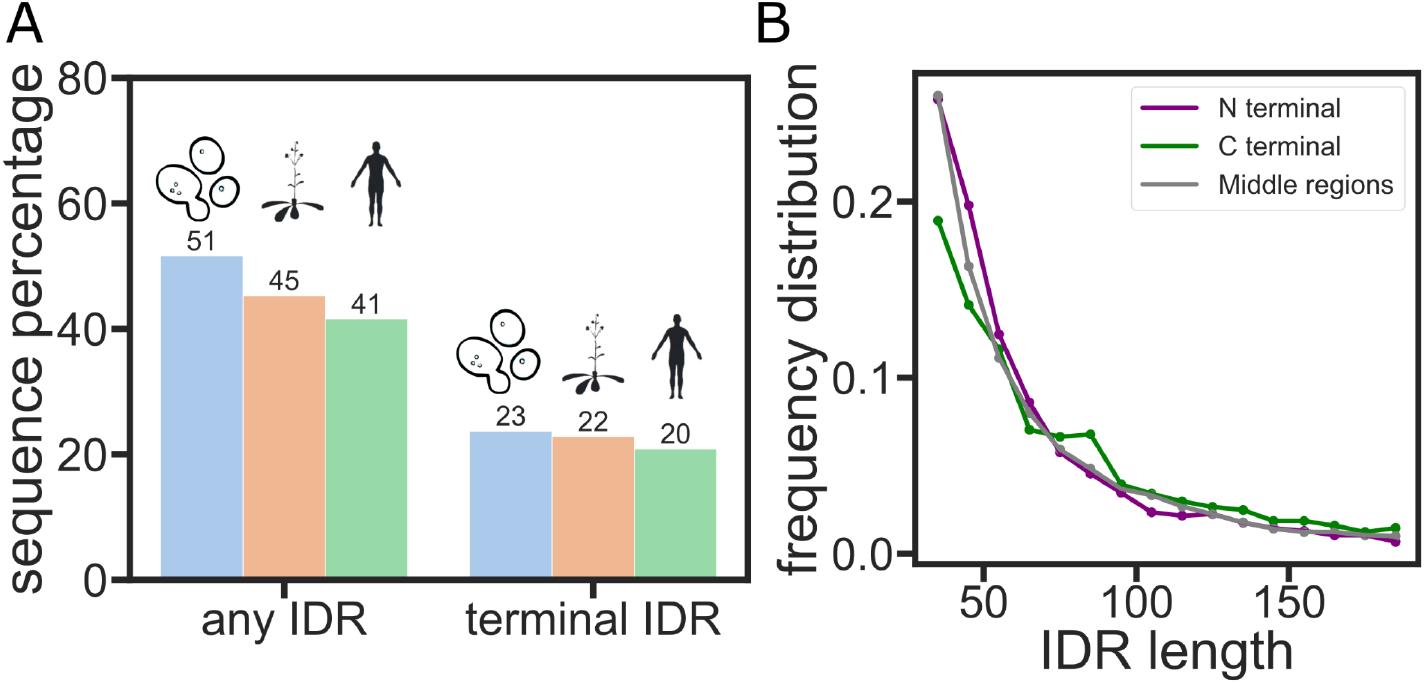
Entropic force may be a widely existing IDR function mechanism in the proteome. **(A)** The percentage of proteins that have a terminal IDR in the yeast, arabidopsis, or human proteomes. **(B)** Distribution of the number of amino acids in the IDRs of the human proteome.

Based on past work, we reasoned that length is a factor that contributes strongly to the entropic force mechanism in these IDRs^9,14^. We therefore wanted to test if there is a significant difference in the length distribution of terminal vs. non-terminal IDRs^45,46^. Our analysis reveals that the length distribution is roughly the same between the terminal and non-terminal IDRs (**Fig. 2B**).

### An IDR simulation database reveals structural diversity

With IDR sequence length being roughly the same in both terminal and non-terminal sequences, we turned our attention to the structural preferences of their ensemble. Ensemble average end-to-end distance (*R*_*ee*_) has been widely used to quantify the global dimensions and the internal structure of dynamic IDR conformational ensembles^19,23^. Since ensemble dimensions cannot be accurately predicted from the sequence, we used the ABSINTH forcefield to gain an atomic-level simulation of over 90 IDR ensembles. Most of these sequences are experimentally validated IDR sequences from the DisProt database^47^ (**Table S1**). These sequences have a diverse distribution of properties including the length, fraction of charged residues (FCR), and net charge per residue (NCPR) (**Fig. 3A-C**).

**Figure 3.**
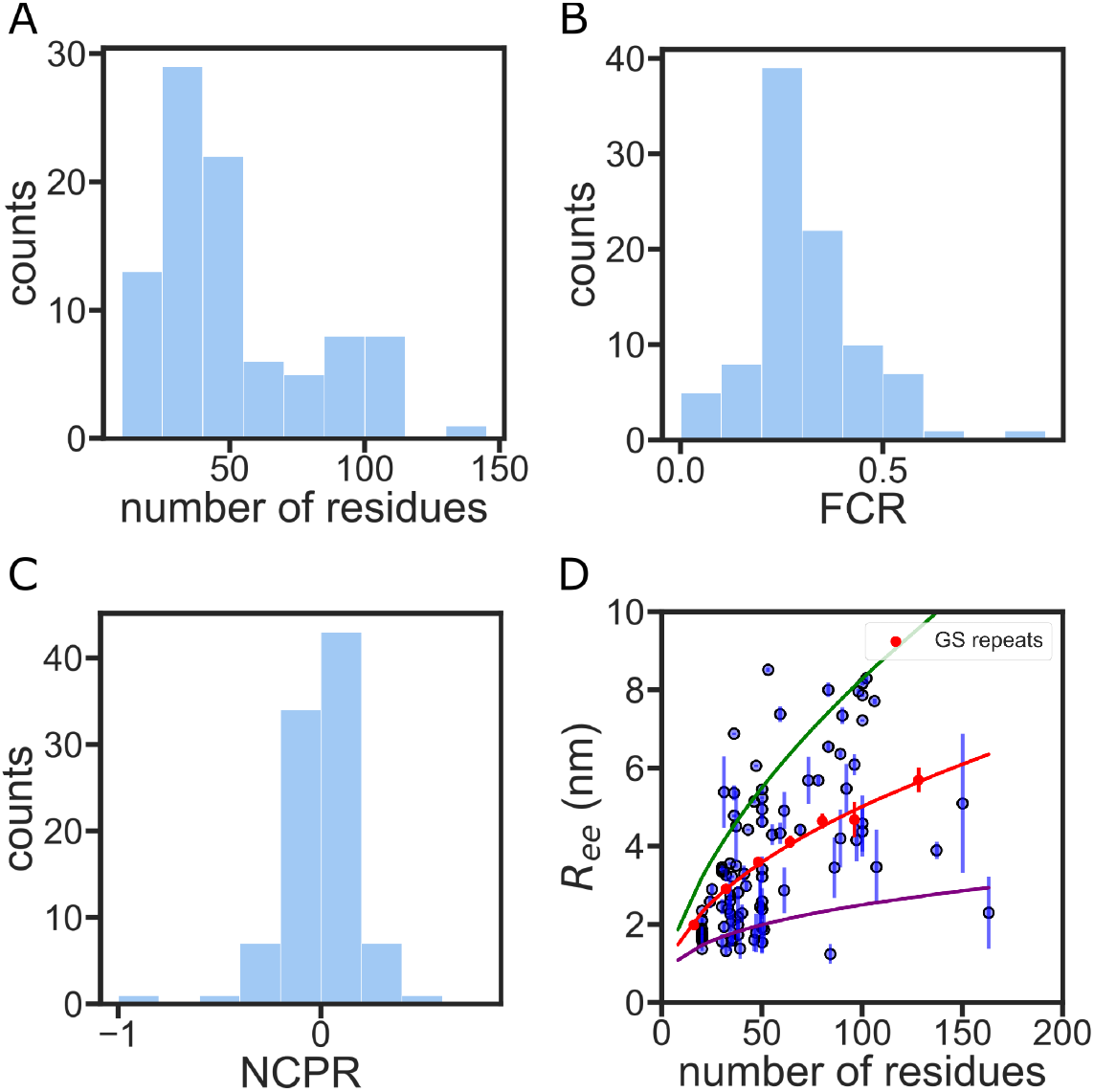
IDR simulation database shows diverse sequence properties and structural preferences. **(A)** The sequence length distribution of the IDR simulation database. **(B)** The fraction of charged residues (FCR) distribution of the IDR simulation database. **(C)** The net charge per residue (NCPR) distribution of the IDR simulation database. **(D)** End-to-end distance vs the number of residues for each simulated IDR. Error bars are calculated from five independent simulations of the same sequence. GS-repeat simulations are shown in red. The red curve is a power law fit of the GS-repeat data. The green curve is the *R*_*ee*_ prediction of the GS-repeats with an exponent of 0.59 and the purple curve is the prediction of the GS-repeats with an exponent of 0.33, which represent the limits of an extended and compact homopolymer^49^.

Simulations reveal a large distribution of *R*_*ee*_ (sometimes more than a factor of 2 for sequences with the same number of amino acids), indicating distinct structural preferences in these sequences. To compare different IDRs of various lengths across the proteome, we use Gly-Ser repeat peptides (GS-repeats) as a homopolymer point-of-reference. It has been shown experimentally that GS-repeats have a similar ensemble to an ideal homopolymer (apolymer where *R*_*ee*_ scales as *N*^0.5^) ^23,48,49^. We simulated several different lengths of GS-repeat sequences using the ABSINTH forcefield. Our simulation data shows *R*_*ee*_ of GS-repeats follows a scaling law with an exponent of 0.48 ± 0.03 (**Fig. S1**), which matches previously reported experimental results^33^. Our analysis shows that a large majority of the sequences measured deviate from the GS-repeat line (**Fig. 3D**).

### Quantifying the entropic force of disordered ensembles using enhanced sampling

We next wanted to probe if these structural preferences alter the magnitude of the entropic force these sequences exert. To assess how ensemble structural preferences change the entropic force, we quantified the change in IDR conformational entropy upon tethering the simulated ensemble to a flat surface and the change in allowed conformations/accessible states (as described in **Methods** and in **Eq. 3**). The change in conformational entropy upon tethering, Δ*S*/*k*_*B*_, is directly correlated to the magnitude of the entropic force (**Fig. 1B**).

To obtain the number of allowed conformations in the tethered state, Ω_*T*_, we tethered our simulated IDR conformational ensemble to a flat surface through the N-terminal *C*_*a*_. Beyond the conformations included in the ensemble, the geometry of the tethering point can also affect the magnitude of Δ*S*/*k*_*B*_. To account for this, we introduced an enhanced sampling method to vary tethering configurations and measure the entropic force at various ensemble orientations relative to the tethered surface (**Fig. 1C, Methods**). With these variations, we generate additional conformations and plot an accessible state heatmap to visualize the number of allowed conformations in each orientation (**Fig. 4A**). To obtain a measure of the entropic force that will be comparable between all sequences, we sum the number of allowed conformations in all different orientations to provide a single entropic force strength for each sequence (**Fig. 4B**).

**Figure 4.**
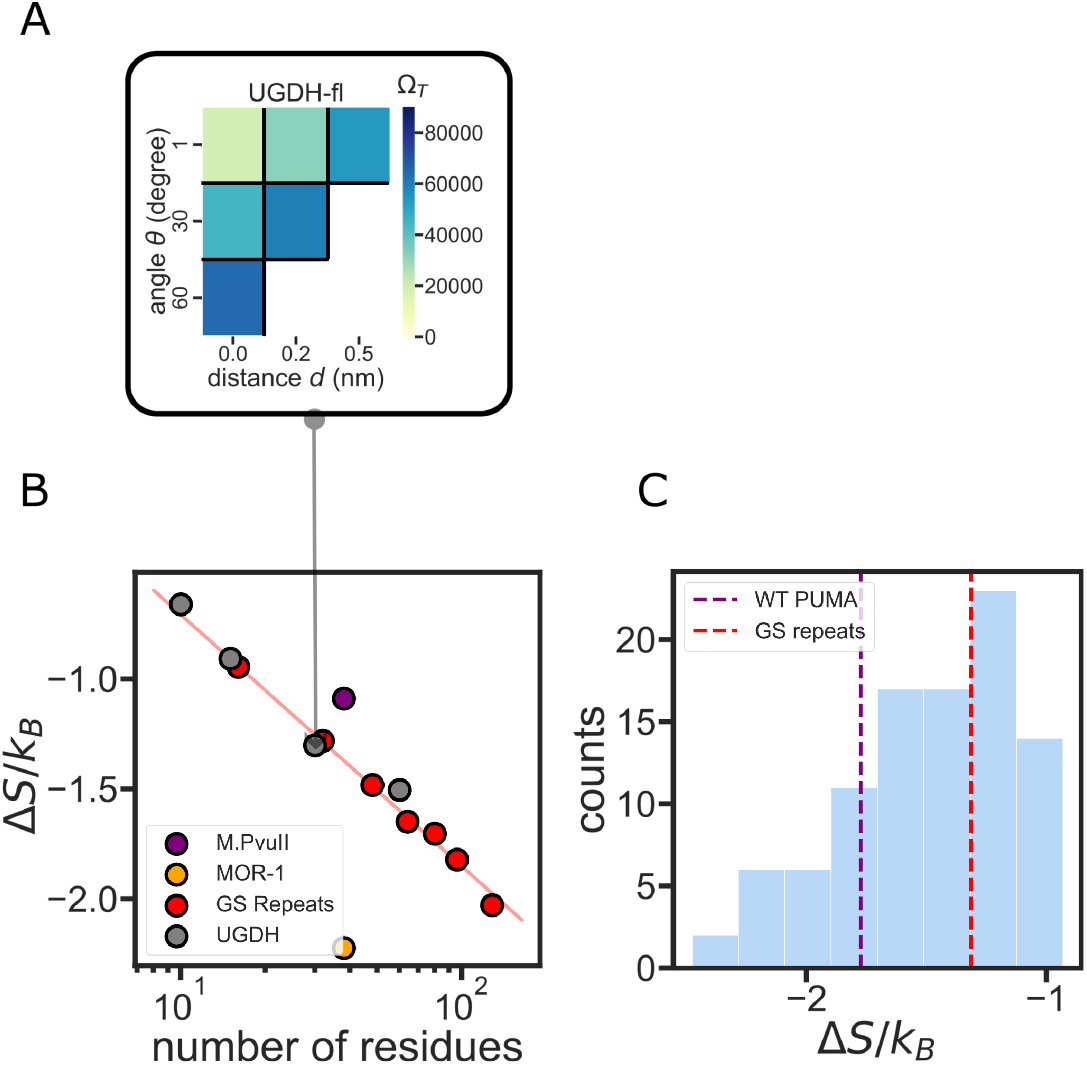
The role of IDR sequence length in determining entropic force strength. (**A**) The variables *d* and θ are varied discreetly to assess the number of allowed states Ω_*T*_ for the ensemble when tethered to the constraint surface. The color in each position on the grid represents the number of allowed states Ω_*T*_ from 6 different ϕ values. The total number of accessible states is used to calculate the entropic force strength for each construct. (**B**) Sequence length determines the entropic force strength of homopolymer-like IDRs. Red curve: an exponential fit of the GS-repeats entropic force strength. Grey dots: UGDH segments as measured in Ref. 14 show a similar entropic force as the equivalent GS-repeat homopolymer. (**C**) A histogram of the entropic force of 96 PUMA scramble sequences. The red dashed line shows the entropic force strength of the same-length GS-repeat sequence.

### Validation of the entropic force calculation using experimental data

Several studies have highlighted the importance of IDR length on the entropic force it exerts^10,12^. A recent study by Keul et al. demonstrated that the length of a terminal IDR tail was the only factor determining its functional effect on the folded enzyme to which it was tethered^14^. The study focused on the C-terminal IDR of a key glycolytic enzyme, UDP glucose 6-dehydrogenase (UGDH). The study showed that the C-terminal IDR acts, through the entropic force it exerts, as an allosteric switch that alters the affinity of the protein to its allosteric feedback inhibitor UDP-xylose. The authors discovered that the entropic force (and the measured binding affinity) depend solely on the length of the terminal IDR, and not on its amino acid composition or sequence. (**Table S1**). As a test of our method, we wanted to see if this length-dependent behavior for the UGDH IDR sequence is reproduced in our simulations.

The homopolymeric GS-repeat entropic force was fitted to an exponential decay function, indicating it is solely determined by the sequence length ^6^. In agreement with Kuel et al.’s observations, UGDH-derived sequences of different lengths also fell on the same line as the GS-repeats (**Fig. 4B**). This indicates that the terminal UGDH IDR has entropic force strength similar to that of a homopolymer. However, UGDH might be a special case resulting from the specific amino acid composition. Indeed, two other IDR sequences display significantly different Δ*S*/*k*_*B*_ despite having the same number of residues (**Fig. 4B**). For example, we selected a disordered region of the type II methyltransferase (M.PvuII, Disprot ID: DP00060r010) from the DisProt database, and compared it to the C-terminal intracellular region of the mu-type opioid receptor (MOR-1, Disprot ID: DP00974r002). Both sequences are 38 residues long. Despite this, the C-terminal region of the MOR-1 has half as many accessible states as M.Pvull when tethered to a constraining surface, generating a stronger entropic force (**Fig. 4B**).

Is the magnitude of the entropic force dependent on amino acid composition alone, or on the sequence of the IDR? To answer this question, we generated a library of scrambled sequences of a naturally occurring sequence, the BH3 IDR domain of the p53-upregulated modulator of apoptosis (PUMA)^50^ (**Fig. 4C**). Despite having the same sequence length and same amino acid composition, scrambles of the PUMA sequence demonstrated a significant difference in entropic force strength. The maximum entropic force of PUMA scrambles is more than two times the minimum force. We observed that scrambled sequences can exert both a stronger and a weaker entropic force upon tethering compared to the wild-type sequence. This result suggests that the order of amino acids in an IDR sequence, and not just amino acid composition, plays a vital role in determining entropic force strength (**Fig. 4C**).

Overall, our simulations recapitulated experimental observables which implicate IDR length as a key factor affecting IDR entropic force, but also highlighted the role of amino acid composition and sequence in the magnitude of this force.

### Systematic analysis of IDR entropic force

We next wanted to understand the role that sequence plays in determining IDR entropic force. We looked for sequence feature correlations with entropic force but found no strong correlations with any individual sequence features (**Fig. S2**). We therefore focused our attention to IDR ensemble dimensions, which are encoded in the sequence but are difficult to predict from structure.^3,19^. We applied our enhanced sampling analysis to 94 IDR sequences obtained from the DisProt database. We observed IDRs generating both higher and lower entropic force compared to GS-repeats, despite having the same length (**Fig. 5A**).

**Figure 5.**
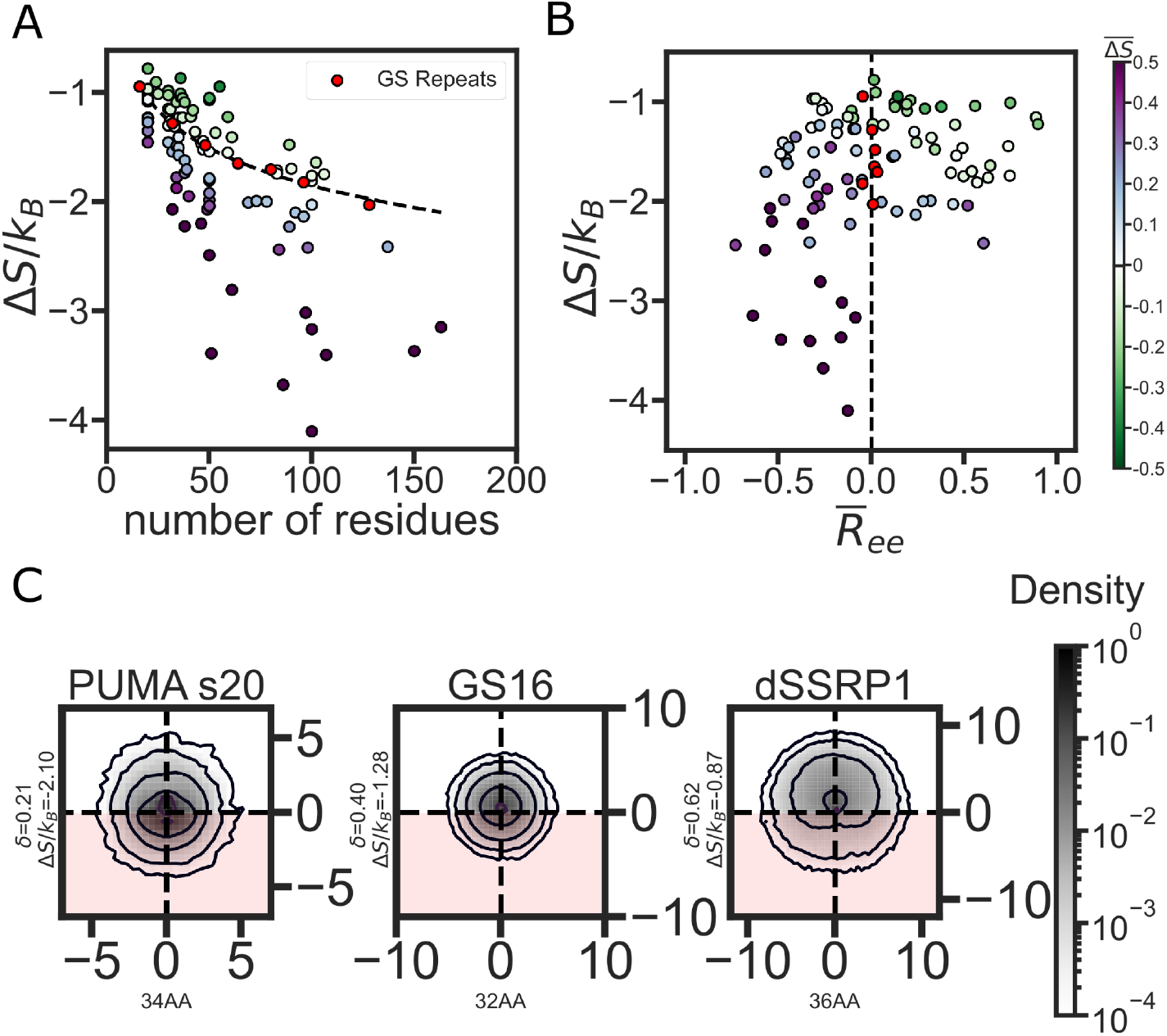
IDR structural preferences divide between weak and strong entropic force. (**A**) Entropic force *vs*. the number of residues in 94 different IDRs. The black curve is an exponential fit of GS-repeat data. Each point represents the entropic force of a single sequence calculated from 5 independent repeats. The color-coding shows the entropic difference between the IDR and the same-length GS-repeat 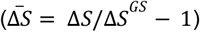, with purple (green) markers showing a stronger (weaker) entropic force compared to the equivalent GS-repeat. (**B**) Entropic force *vs* the GS-repeat normalized end-to-end distance 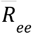 (see **Eq. 3**). Each marker represents a single IDR color-coded as in (A). (**C**) XZ-projections of *C*_α_ density for 3 different IDRs with increasing asphericity. The constraint plane is normal to Z=0 such that the density at Z>0 will avoid the surface and the density at Z<0 clashes with the surface (the disallowed region is indicated by the red color).

To ascertain how ensemble dimensions may play a role in determining Δ*S*/*k*_*B*_, we must first find a way to compare the ensembles of IDRs of various lengths. To do this, we normalize the average *R* of all IDRs against the *R*_*ee*_ of a GS-repeat sequence of the same length to get normalized end to end distance 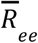 (**Eq.1, Method**). 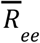 has a negative value when the ensemble is more compact than a GS-repeat, and a positive value when an ensemble is more expanded. We plot Δ*S*/*k*_*B*_ for each sequence as a function of this normalized distance in **Fig. 5B**. It is immediately noticeable that the vertical red line drawn at 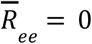 separates sequences with a higher entropic force (purple markers) from those with a weaker entropic force (green markers). This means ensembles that are on average more compact than an equivalent GS-repeat (as indicated by a negative 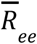) tend to generate a stronger entropic force, while more expanded ensembles tend to generate a weaker entropic force than equivalent GS-repeats.

This seemed counterintuitive since our initial thought was that an expanded ensemble should take up more space and would therefore lose more conformational entropy upon tethering to the constraint surface. However, a more expanded ensemble will tend to have a higher persistence length and a more ellipsoid shape^51^. These properties mean that the backbone will point away from the tethered surface (because of this longer persistence length), reducing the number of conformations that will sterically clash with the surface. To validate this hypothesis, we calculated the average asphericity of the IDR ensemble^37^.

Similar to 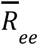, ensembles with low asphericity have a lower entropic force, and ensembles with a high asphericity have a stronger entropic force than that of GS-repeats (**Fig. S3, S4**). This suggests that a more spherical ensemble tends to have a higher possibility of clashing with the constraining surface and thus generates a stronger entropic force, while a more elongated ensemble tends to have less interaction with the constraining surface. To verify this, we visualized the position of *C*_α_ atoms on an XZ plane that is normal to the constraining surface for several sequences (**Fig. 5C**). This visualization highlights how spherical ensembles with a low asphericity tend to have more atoms at or under the constraint surface (located at Z=0) while ellipsoidal ensembles with a high asphericity tend to expand with a higher atom density above the constraint surface.

### Changes in solution chemistry alter IDR entropic force strength

An alternative way to change ensemble dimensions, and one that does not involve a change in IDR sequence is to expose IDRs to different solution environments^16,40^. Previously, we found that IDRs tend to be more sensitive than folded proteins to changes in the chemical composition of their surrounding solution. We designed the Solution Space Scanning method to simulate IDR ensemble structural preferences under changing solution conditions^40^. Briefly, the method alters IDR ensembles by tuning the protein backbone:solution interactions of the ABSINTH forcefield to be more or less repulsive than the value for water (see **Methods**). Usually, IDRs have a more compact conformational ensemble in repulsive solutions (e.g. in the presence of an osmolyte or a more crowded environment). In attractive solutions (e.g. urea or other denaturants), IDRs have an expanded conformational ensemble. However, this general trend can be mitigated and sometimes even reversed based on the IDR sequence^16,17,40^.

To see how solution-induced changes in the ensemble affect entropic force, we used Solution Space Scanning to simulate the ensemble average *R*_*ee*_ of the proteins shown in **Fig. 2B** and **Fig. 5** in five different solution conditions. We observed significant compaction of the ensemble in the repulsive solution, and the ensemble change is correlated with protein:solvent interaction strength (**Fig. 6A**). To quantify how Δ*S* changes with solution condition change, we use the change in entropic force between solute and buffer with the following equation:

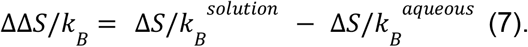

**Figure 6.**
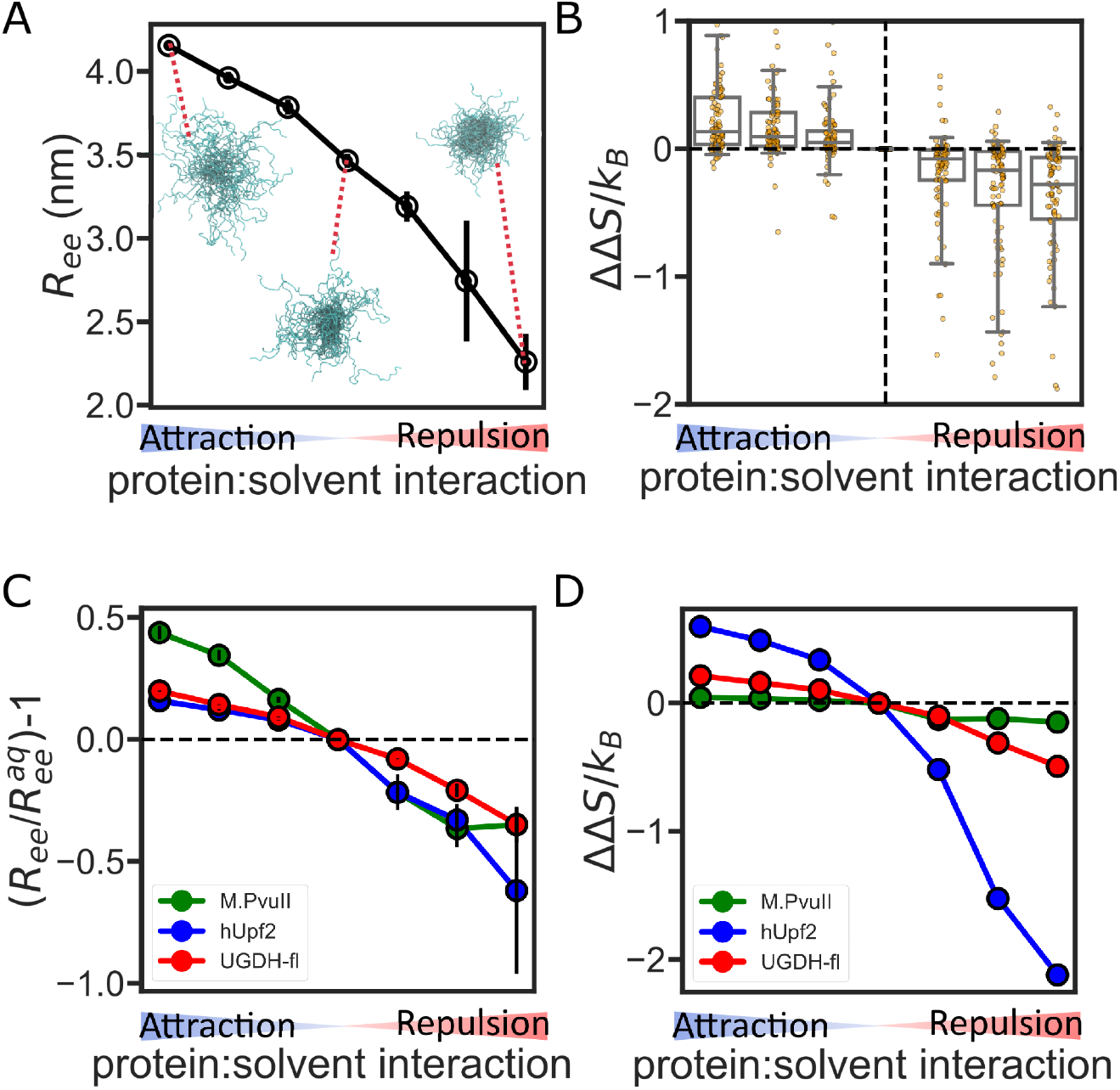
Solution conditions alter IDR entropic force. (**A**) End-to-end distance for UGDH-fl as a function of backbone:solution interactions. The blow-up ensembles show representative conformations in attractive, neutral (aqueous), and repulsive solutions. (**B**) Box plot showing the change in entropic force due to change in protein backbone:solvent interactions. Boxes show the median as a central line, the median 50% as the box limits, and the median 90% of the data as the whiskers. Individual sequences are shown as points overlaid on each box. **(C)** Solution sensitivity of three IDR ensembles. Solution sensitivity is quantified using relative *R*_*ee*_ compared to the *R*_*ee*_ of the same IDR in the neutral (aqueous) solution. **(D)** The change in entropic force due to solution condition changes for the three IDR ensembles.

Here,ΔΔ*S*/*k*_*B*_ represents the change in the entropic force in different protein:solution interactions. We calculate the entropic force change between the buffer/aqueous condition and other solution conditions. Our analysis shows that, on average, IDRs will generate a stronger entropic force when their ensemble is compacted due to the presence of a repulsive solution (**Fig. 6B**). This result strengthens our conclusion that compact IDR ensembles tend to exert a larger entropic force than extended ensembles.

However, not every IDR is sensitive to solution condition changes. We observed that some IDRs do not have a significant entropic force change, despite significant changes in their ensemble (**Fig. 6C, D**). For example, M.PvuII displays a significant change in *R*_*ee*_, but almost no change in entropic force (**Fig. 6C**). On the other hand, the ensemble of the IDR of the regulator of nonsense transcripts 2 (hUpf2, Disprot ID: DP00949r013) is very sensitive to solution changes, and Δ*S*/*k*_*B*_ changes accordingly. This suggests that solution-driven changes in entropic force response are highly sequence-dependent. Different sequences encode diverse structural ensembles that in turn influence IDR environmental response. Interestingly, the UGDH IDR has a low sensitivity of entropic force despite its high sensitivity ensemble. Considering UGDH performs an allosteric function through its entropic force, this suggests some sequences may evolve to generate a stable entropic force for performing their function.

## Conclusions

Here we report on a computational method to quantify the conformational entropic force of tethered IDRs using all-atom Monte Carlo simulations. Compared to coarse-grained or analytical models, this method offers an accurate, quantitative metric of how IDR entropic force is determined by sequence-encoded conformational ensemble preferences. Our method is compared against and qualitatively matches previously published experimental measurements of entropic force (**Fig. 4B**). Our results also support the current literature and highlight that IDR sequence length is indeed a key factor in the entropic force it exerts (**Fig. 4B**). Despite its drawbacks and limitations (see the section in **Methods**), our method offers an accessible description of the entropic force which is computationally easy to calculate and a self-consistent dataset from which to draw conclusions linking between IDR sequence and entropic force.

Our simulations show that there is more to the story of entropic force than just the length of the sequence. We reveal that IDR structural preferences can determine the magnitude of entropic force strength. We show that the structural preferences of IDR ensembles are encoded not just in amino acid composition but also in their arrangement in the sequence, which can be an important factor in determining entropic force strength. Perhaps counterintuitively, we find that more expanded IDR ensembles can extract a weaker entropic force than more compact IDR ensembles when tethered to a flat surface (**Fig. 5A, 5B, S4**).

We also show that the entropic force exerted by an IDR can change when the surrounding chemical environment changes. By modulating protein backbone:solvent interactions, we altered IDR ensembles and showed that the entropic force magnitude of most IDRs increased as their ensembles became more compact, validating the trend shown for different sequences (**Fig. 6B**). This result also suggests the possibility of manipulating IDR entropic force by altering the physical-chemical composition of the cellular environment ^16,17,52^.

Since the dimensional properties of IDR sequences are sequence-encoded^53^, we propose that some sequences have evolved to exert an outsized entropic force on the protein they are tethered to, while other sequences have evolved to exert a weak force. Our study further suggests that this entropic force can be modulated by post-translational modifications and changes in the cellular environment that are known to alter IDR ensembles^1,2,17,54^. Taken together, the entropic force is a sequence-encoded, tunable function that may be more common than previously realized in IDR-containing proteins.

## Supporting information

Supplementary text contains Figures S1-S4.

Table S1 contains all IDRs sequences, the ensemble analysis result, and the entropic force analysis result of the simulation database.

## Supplementary information

Supplementary text contains **Figures S1-S4**.

**Table S1** contains all IDRs sequences, the ensemble analysis result, and the entropic force analysis result of the simulation database.

All code used in this manuscript is provided at: https://github.com/sukeniklab/Entropic_Force

## Acknowledgments

We thank the NSF-CREST: Center for Cellular and Biomolecular Machines at UC Merced (NSF-HRD-1547848 and NSF-HRD-2112675) for the computational fellowship awarded to FY. We thank David Moses for his comments and suggestions. Research reported in this publication was supported by the NIH under award R35GM137926 and the NSF under IntBio-2128067 to SS. We acknowledge computing time on the MERCED cluster at UC Merced, NSF Grant ACI-1429783, and on the XSEDE computational infrastructure framework, Grant No.TG-MCB190103 to SS, supported by NSF Grant ACI-154856, and on the Expanse cluster through HPC@UC, supported by NSF Grant OAC-1928224.

