## Supplementary text contains Figures S1-S4. for "Structural preferences shape the entropic force of disordered protein ensembles"

Feng Yu[1] and Shahar Sukenik[1,2]

1. Quantitative Systems Biology Program, University of California, Merced, California, United States
2. Department of Chemistry and Biochemistry, University of California, Merced, California, United States

**Figure S1**

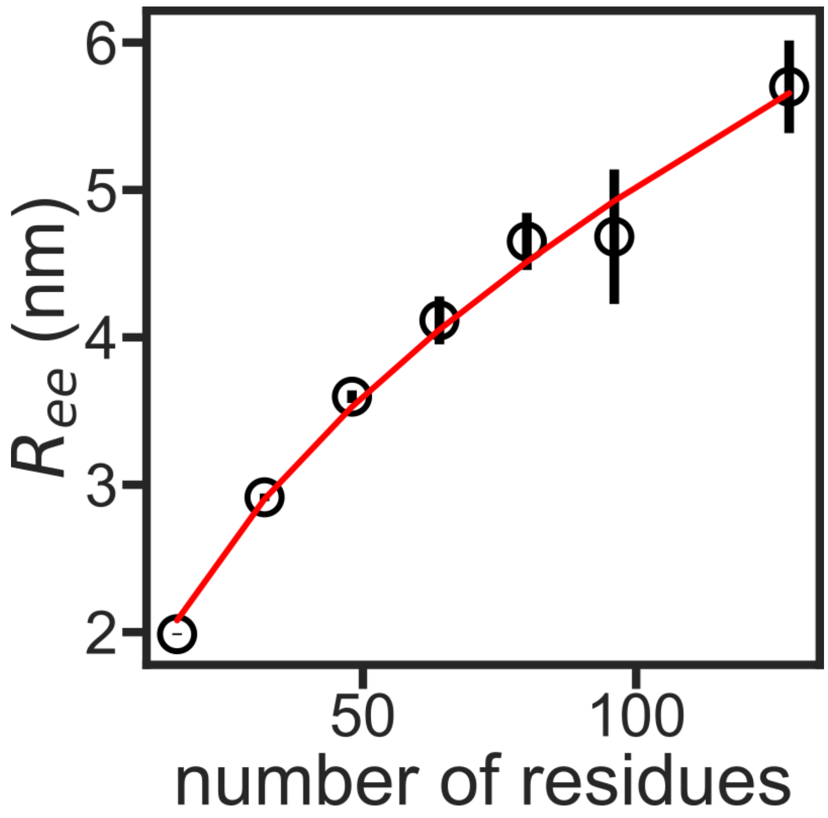


**Figure S1. GS repeats match homopolymer scaling law under buffer conditions.** The average end-to-end distance from five repeats vs the total number of residues for a series of Gly-Ser repeats, The error bars are the standard deviation of the five repeats. The red curve is the result of fitting to $R_{ee} = R_0 N^{\nu}$, with $R_0 = 0.55 \pm 0.06$ nm and $\nu = 0.48 \pm 0.03$. Errors are obtained from the fit.

**Figure S2**

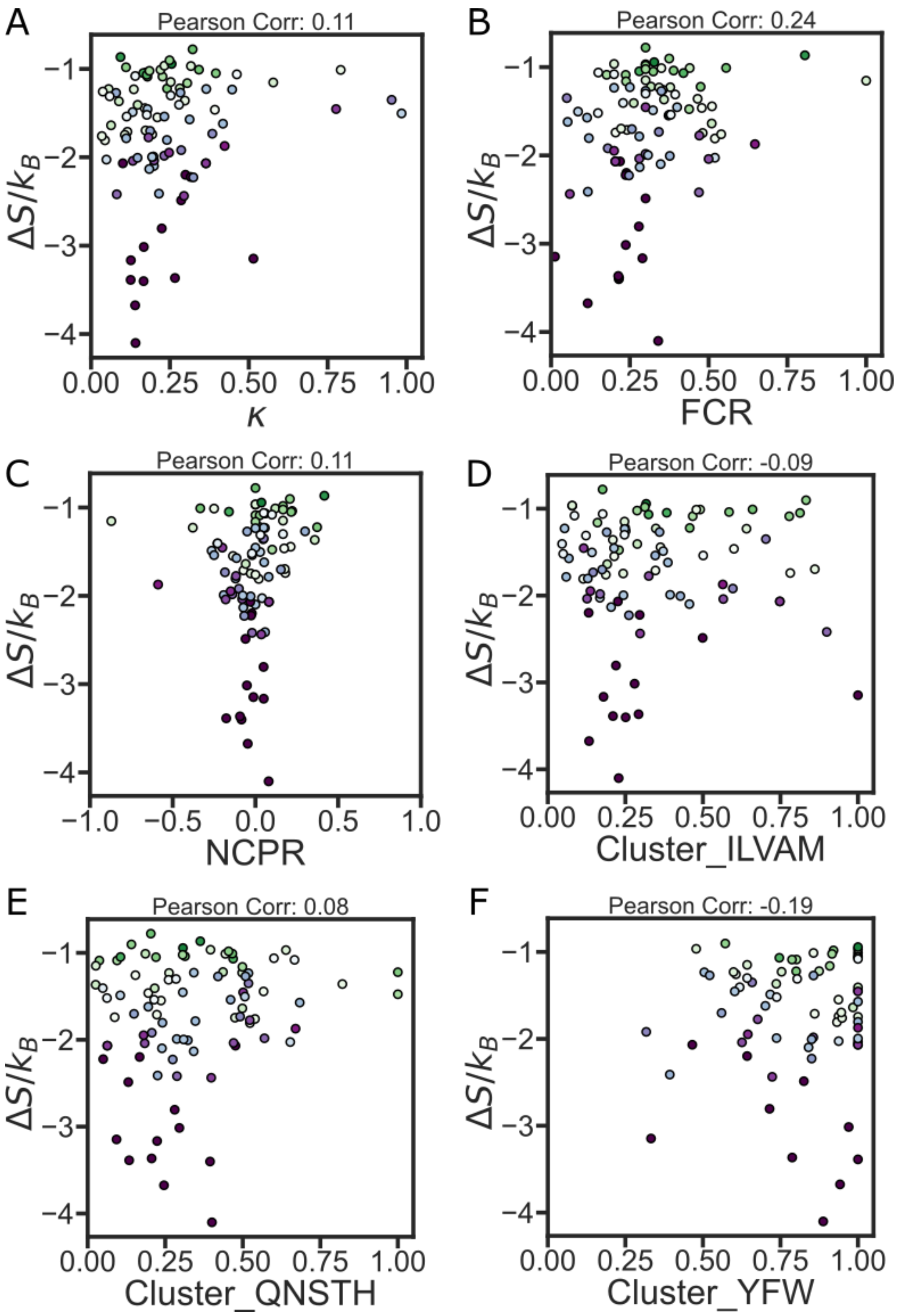


**Figure S2. Sequence features are not correlated with the entropic force strength.** The entropic force strength is plotted vs several sequence features calculated using the localCIDER python package. (**A**) κ: a metric for mixing of charged amino acids[19], (**B**) FCR: fraction of charged residues, (**C**) NPCR: net charge per residue, (**D**)Cluster_ILVAM: hydrophobic amino acid mixing calculated using the same algorithm as κ, (**E**) Cluster_QNSTH: polar amino acid mixing calculated using the same algorithm as κ, (**F**) Cluster_YFW, aromatic amino acid mixing calculated using the same algorithm as κ.

**Figure S3**

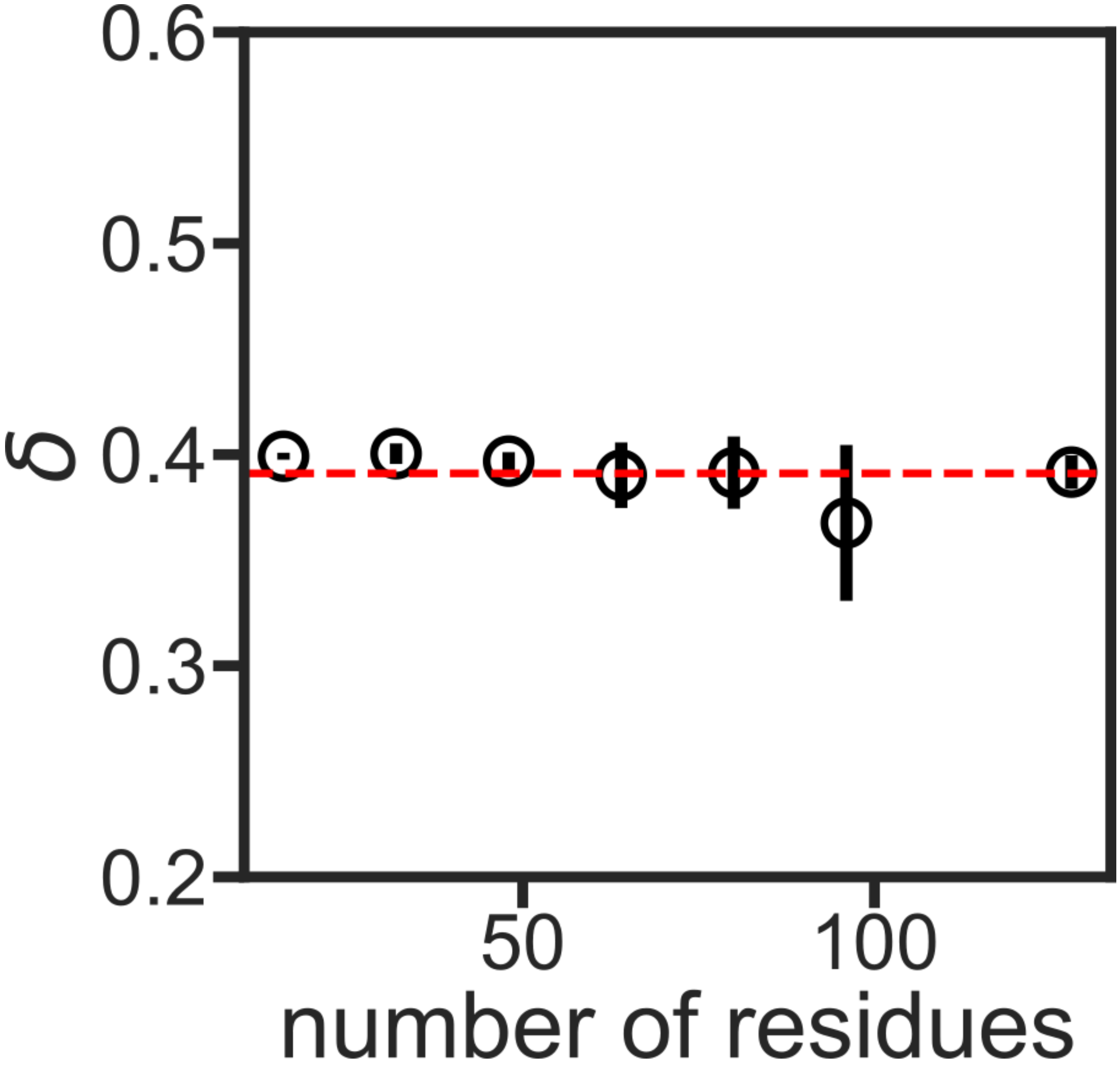


**Figure S3. GS repeat asphericity is independent of length.** The average asphericity of GS repeats vs the number of residues in the sequence. The mean of all seven data points is shown by the red line, with $\delta = 0.39 \pm 0.01$.

**Figure S4**

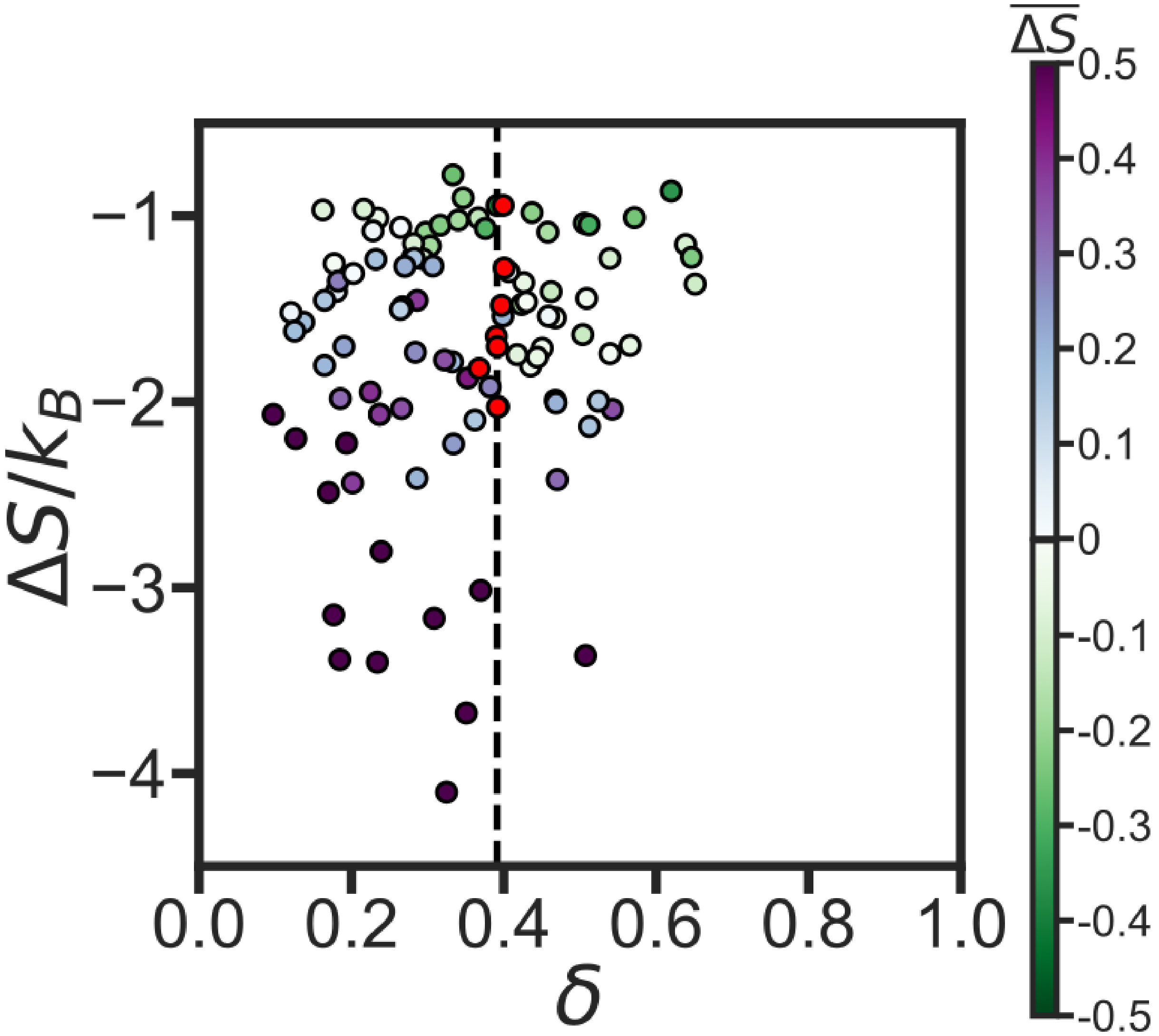


**Figure S4.** Entropic force as a function of average asphericity. The black line represents the length-independent asphericity of GS-repeats shown in **Fig. S3**. Each marker represents a single IDR, color-coded as in **Fig. 5A** , with stronger purple (green) markers showing a stronger (weaker) entropic force compared to the GS repeat of the same size.
